# Dashing Growth Curves – a web application for rapid and interactive analysis of microbial growth curves

**DOI:** 10.1101/2022.12.16.520708

**Authors:** Michael A. Reiter, Julia A. Vorholt

## Abstract

**Background:** Recording and analyzing microbial growth is a routine task in the life sciences. Microplate readers that record dozens to hundreds of growth curves simultaneously are increasingly used for this task, raising the demand for their rapid and reliable analysis.

**Results:** Here, we present Dashing Growth Curves, an interactive web application (http://dashing-growth-curves.ethz.ch/) that enables researchers to quickly visualize and analyze growth curves without coding knowledge and independent of operating system. Growth curves can be fitted with Logistic or Gompertz growth models, or manually by selecting the linear growth phase in a logarithmic plot. Furthermore, Dashing Growth Curves automatically groups replicate samples and generates downloadable plots for extracted growth parameters such as growth rate or lag time.

**Conclusions:** Dashing Growth Curves is an open-source web application that reduces the time required to analyze microbial growth curves from hours to minutes.

## Background

Recording of growth curves is an integral part of microbial research into single celled organisms. Monitoring microbial abundance over time allows researchers to determine growth parameters such as lag time, maximum growth rate, or maximum population size among others under specific conditions. In turn, these can be used, for example, to compare different organism, to investigate the effect of media formulations, to quantify the impact of concentrations of growth inhibitory compounds and the effects of temperature or oxygen availability. Over the last decades, microplate readers that can record many samples in parallel have dramatically increased the amount of data an individual researcher can generate. While it used to be a couple dozen samples, it is now routinely hundreds to thousands in one experiment.

Growth curves can be analyzed manually in programs such as Microsoft Excel. However, this is slow and error prone. To alleviate this problem, several tools have been developed to aid in the analysis. However, most require coding skills, such as GrowthCurver^1^, Grofit^2^ or growthrates^3^ and some are closed-source commercial software (e.g. GrowthRates^4^). To streamline the process of growth curve analysis and make it more accessible, we developed Dashing Growth Curves, an easy-to-use open-source web application.

## Implementation

Dashing Growth Curves is written in Python (3.11) and uses the Plotly Dash (2.7.0) framework (https://dash.plotly.com/) for the web application logic. Graphs are plotted with Plotly (5.11.0) (https://plot.ly). Data handling is done with pandas (1.5.2)^7^. Numerical calculations and curve fitting are done with Numpy (1.23.5)^8^, Scipy (1.9.3)^9^ and Uncertainties (3.1.7)^10^. Compute intensive tasks (e.g. curve fitting) are handled on a separate task queue using Celery (5.2.7) (https://docs.celeryq.dev/en/stable/reference/index.html) and Redis (4.3.5) (https://redis.io/). User interface icons are from the Bootstrap icon library (https://getbootstrap.com/). The source code is available on GitHub (https://github.com/mretier/growthdash).

### Fitting growth curves

Dashing Growth Curves makes the Logistic and Gompertz growth models as modified by Zwietering and co-workers^5^ available to fit growth data (**Error! Reference source not found**.). The models explicitly contain the maximum growth rate, the lag time and the maximum population size as parameters. The user can choose between the standard definition of the lag time^5^ and a an additional definition of lag time referred to herein as “tight”. The former is the timepoint when the tangent at the inflection point of the fitted sigmoidal curve intersects the x-axis^5^. The latter is defined as the timepoint when the slope of the fitted sigmoidal curve increases most (i.e. the smallest zero of the third derivative of the sigmoidal curve)^6^, which generally occurs later than the standard definition of the lag phase and is closer to the segment of the sigmoidal curve that can be approximated by a line. Analogously, the end of the logarithmic growth phase occurs when the tangent at the inflection point intersects the line *y* = *A* (the maximum population size) or at the timepoint where the third derivative of the sigmoidal function has its largest zero, respectively.

**Table 1:** *y*: predicted logarithmic population size (dependent variable); *t*: time (independent variable); *A*: maximum population size; *µ*_*max*_: maximum growth rate; *λ*: lag time.

| Growth model | Equation |
| --- | --- |
| Modified Logistic | $y = \frac{A}{1 + \exp \left[ \frac{4\mu_{max}}{A} (\lambda - t) + 2 \right]}$ |
| Modified Gompertz | $y = A \cdot \exp \left\{ - \exp \left[ \frac{\mu_{max} \cdot e}{A} (\lambda - t) + 1 \right] \right\}$ |

### Data privacy and local installation

Dashing Growth Curves does not store user data nor uploaded data beyond the time it is in active use in the web application. Data uploads are not logged. Cookies are not set. Additionally, Dashing Growth Curves can be installed locally within a few minutes following a short manual (see the readme in the GitHub repository). The local installation is identical (except for the backend implementation) to the hosted web version.

## Results and Discussion

Dashing Growth Curves can be accessed via the internet using any common web browser (http://dashing-growth-curves.ethz.ch/). However, the application has been optimized for best display in Google Chrome.

On the Dashing Growth Curves landing page users can upload their data and are offered different resources (Figure 1). For Dashing Growth Curves to be able to parse growth data, it needs to be in a simple table format and contain timestamps and sample names (Figure 2). After data upload, it is embedded in a graphical user interface. Two views of the data are available. The “sample view” shows the growth curve of a sample and its associated growth parameters and blanks (Figure 3). The “summary view” plots all computed growth parameters and groups replicate samples (Figure 4). Dashing Growth Curves extracts the doubling time, the maximum growth rate, the lag time, the number of doublings from lowest to highest measured population size, the doublings in logarithmic growth phase and the yield (i.e. the maximum population size) for every sample. If no logarithmic growth phase is observed the associated parameters are left undetermined. By default, the population size is indicated by optical density measurements at 600 nm (OD_600_), but this can be changed in the settings menu (e.g. to colony forming units, Figure 3 ④).

**Figure 1.**
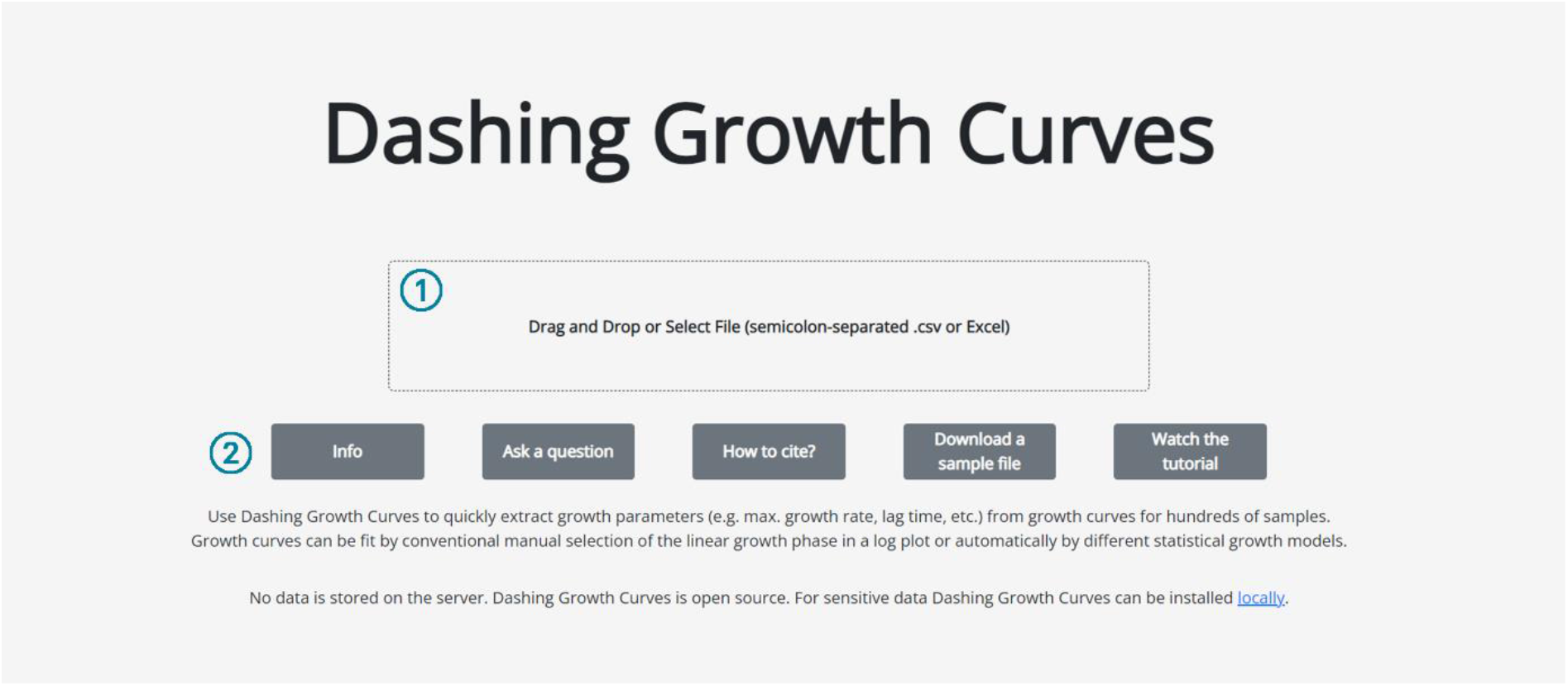
Dashing Growth Curves landing page. ① Drag and drop data file into box to upload it to the application or click inside the box to open a dialog box to select it. ② Different resources. “Info” forwards to the GitHub repository (https://github.com/mretier/growthdash) which contains all information about Dashing Growth Curves. “Ask a question” forwards to the associated GitHub issues page where bugs can be reported or questions about the application can be raised (https://github.com/mretier/growthdash/issues). “How to cite” forwards to the most recent associated publication of Dashing Growth Curves. “Download a sample file” downloads an Excel file with correctly formatted growth curve data. “Watch the tutorial” forwards to a video tutorial explaining Dashing Growth Curves (https://www.youtube.com/watch?v=lhvgZyPlHzA).

**Figure 2.**
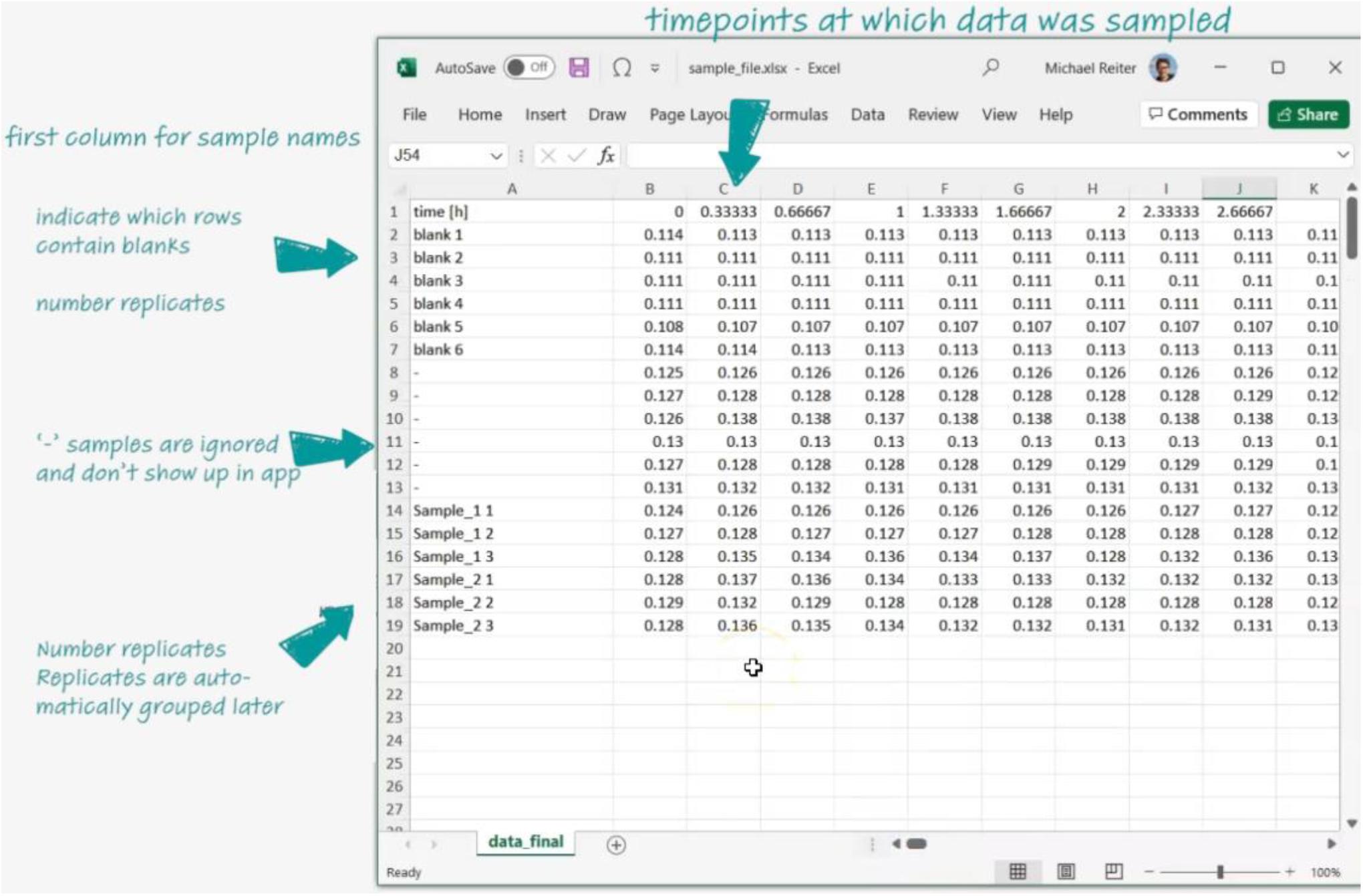
Data structure of a set of individual growth curves.

**Figure 3.**
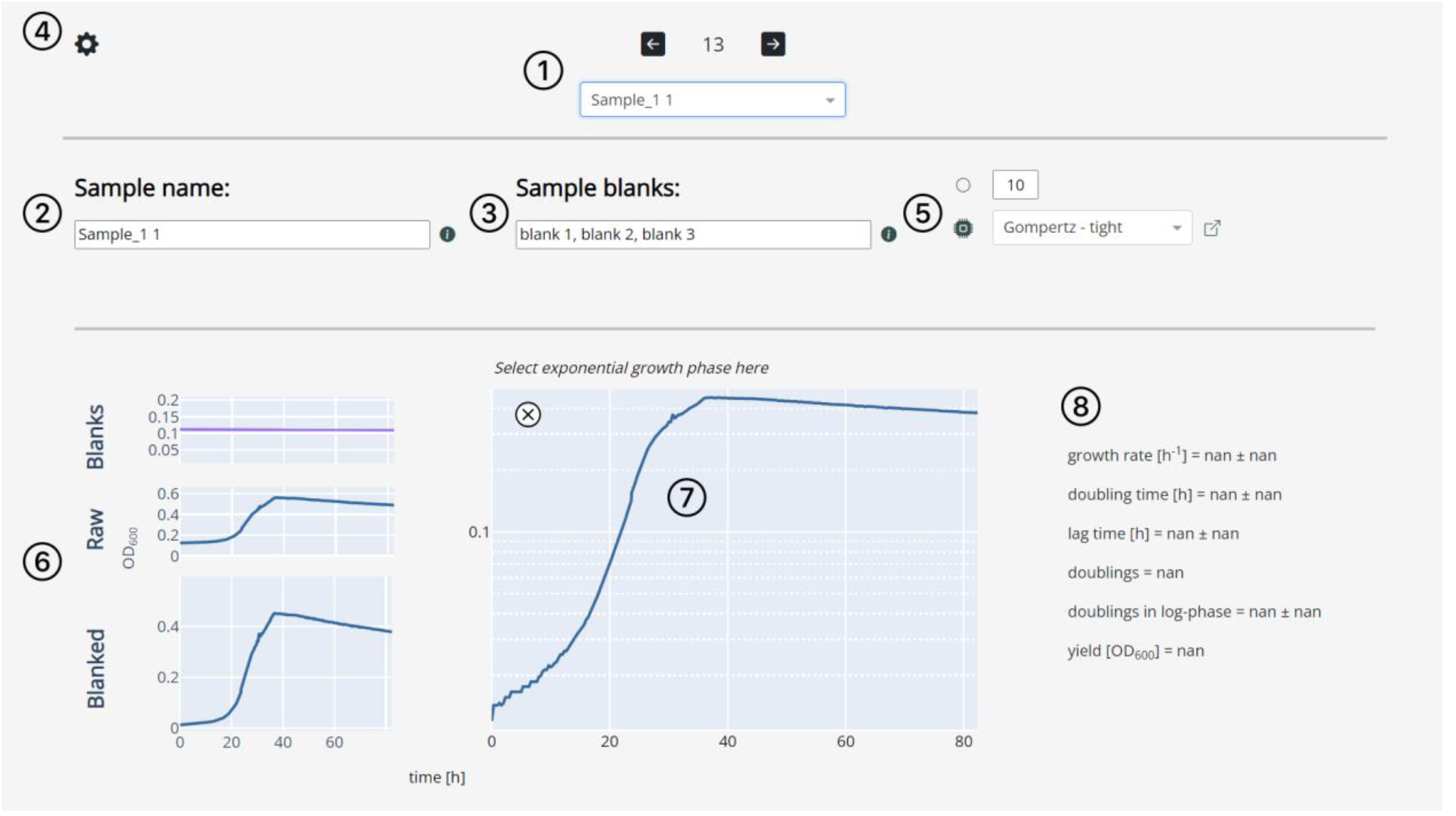
Dashing Growth Curves sample view. ① Position of the currently selected sample and its name. The dropdown menu or the arrow buttons can be used to change the currently displayed sample. If there are 6, 12, 24, 96 or 384 samples, a sample position will be associated with each sample (e.g. the 13^th^ row in a data file of 96 samples will be position B2) if there are a number of samples that are not associated with a standard microtiter plate size, each will be associated with a number. ② The sample name can be changed here. ③ The field displays the blanks associated with the current sample. The associated blanks can be changed by providing a list of sample names that are to be used as blanks. By default, Dashing Growth Curves takes the first three samples in a data file as blanks. The default blanks for all samples can be changed in ④. ⑤ Circle button: data can be smoothed using a rolling average algorithm (the default window is 10 data points). CPU button: Fit all growth curves to a chosen parametric growth model. ⑥ Graphs of the associated blanks and their mean (here all three blanks are very similar and fall onto one line), the raw trace of the currently selected sample and the blanked trace. ⑦ The blanked trace plotted on a logarithmic y-axis. Pressing the ‘x’ button excludes the sample from the summary view (Figure 1 Figure 4). ⑧ Growth parameters of the currently selected sample. If the logarithmic growth phase has not been determined manually or the sample has not been fitted to a parametric growth model yet, the values are ‘nan’ (not a number).

**Figure 4.**
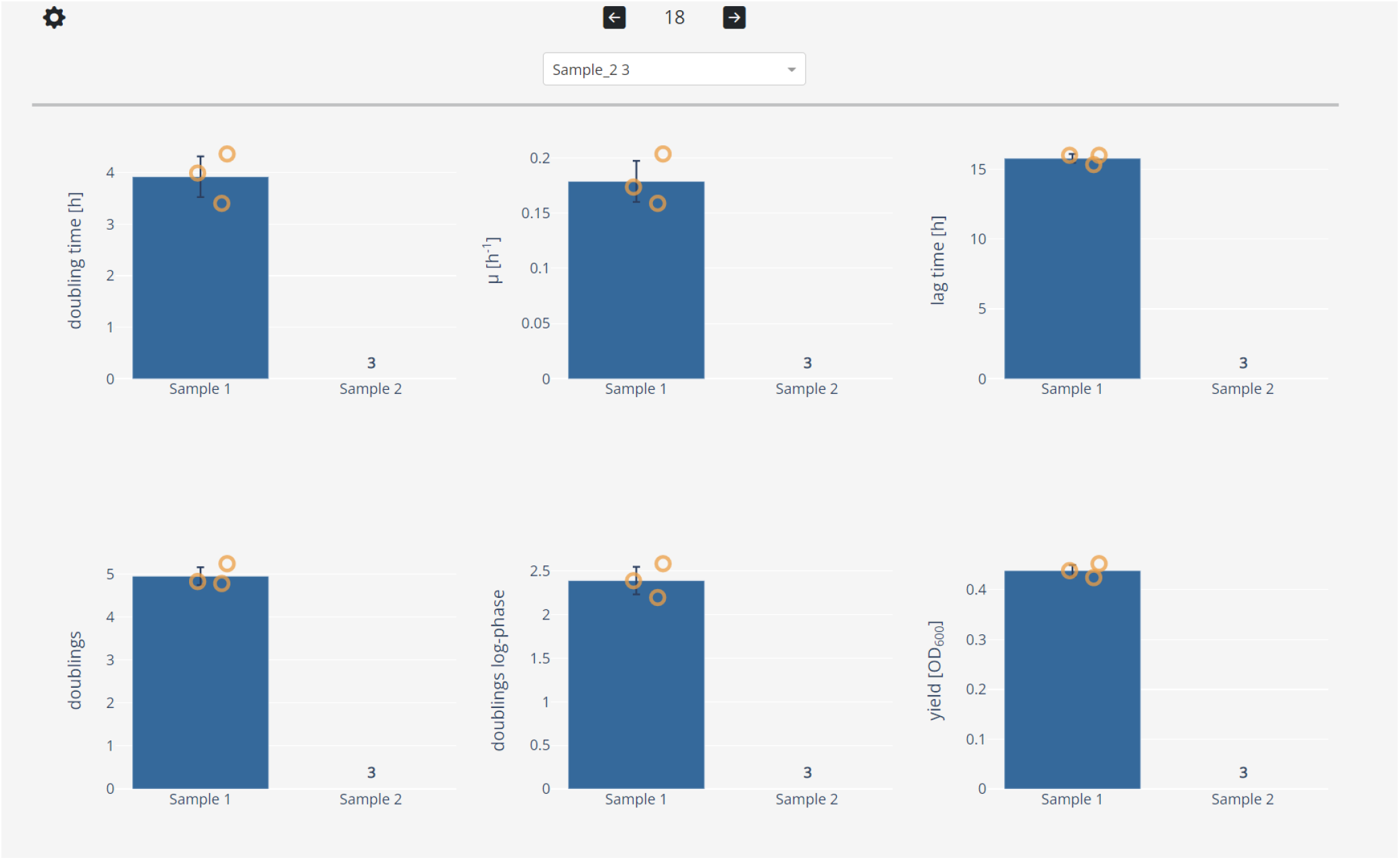
Dashing Growth Curves summary view. Sample replicates are automatically grouped. For each group and every growth statistic a bar chart is plotted on top of which the individual datapoints are overlayed. Clicking on a datapoint selects it and displays its growth curve in the sample view (e.g. to doublecheck outliers). Individual growth curves can be excluded (e.g. for technical reasons). Here, for ‘Sample 2’ all three replicates were excluded. The number of excluded samples is tracked by a counter above the sample name.

To determine growth parameters, different options are available. Data can be fitted automatically with parametric models (Figure 5B and section “Fitting growth curves”), or each growth curve can be analyzed individually by manually selecting the segment of the growth curve that exhibits a linear increase over time in a logarithmic plot and fitting a line (Figure 5C). Manual selection, while not taking into account the whole growth curve, provides greater flexibility for more complex growth patterns (e.g. diauxic growth). In contrast, automatic selection enables faster throughput, but the correct fitting of complex growth profiles may become difficult to impossible.

**Figure 5.**
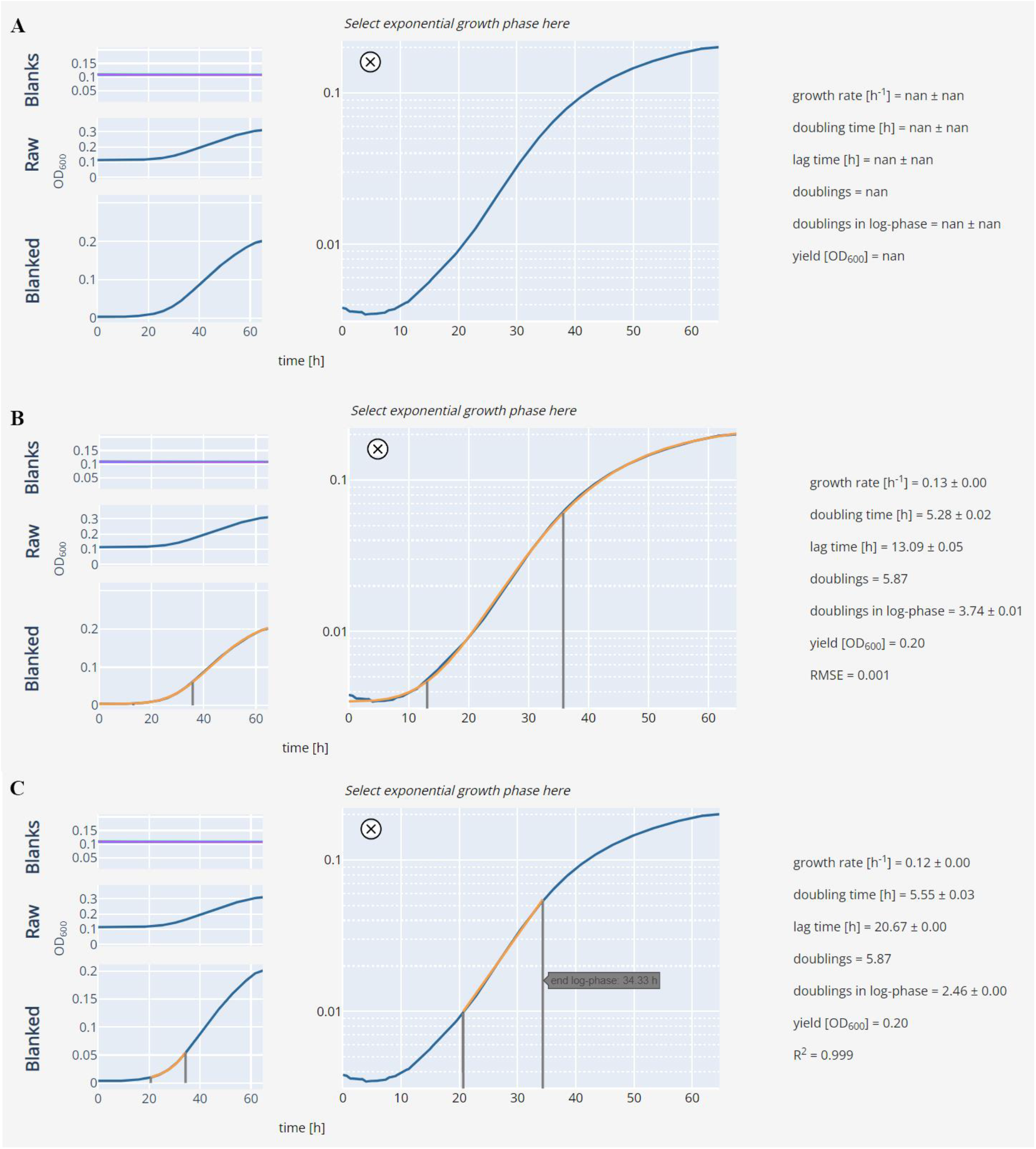
Fitted growth curve examples. **A)** Raw, unfitted growth curve. **B)** Growth curve fitted with a Gompertz model (see “Fitting growth curves” section). The fit is overlayed in orange. The grey lines indicate the beginning and end of the logarithmic growth phase. **C)** Growth curve fitted manually by selecting the segment in which the observed logarithmic growth over time is linear.

To facilitate further analysis and data presentation, all plots can be downloaded as editable vector graphics (.svg files, Figure 6 ①) or interactive HTML plots (Figure 6 ②). Furthermore, a summary file that contains all data can also be downloaded for further analysis or custom plotting (Figure 6 ③).

**Figure 6.**
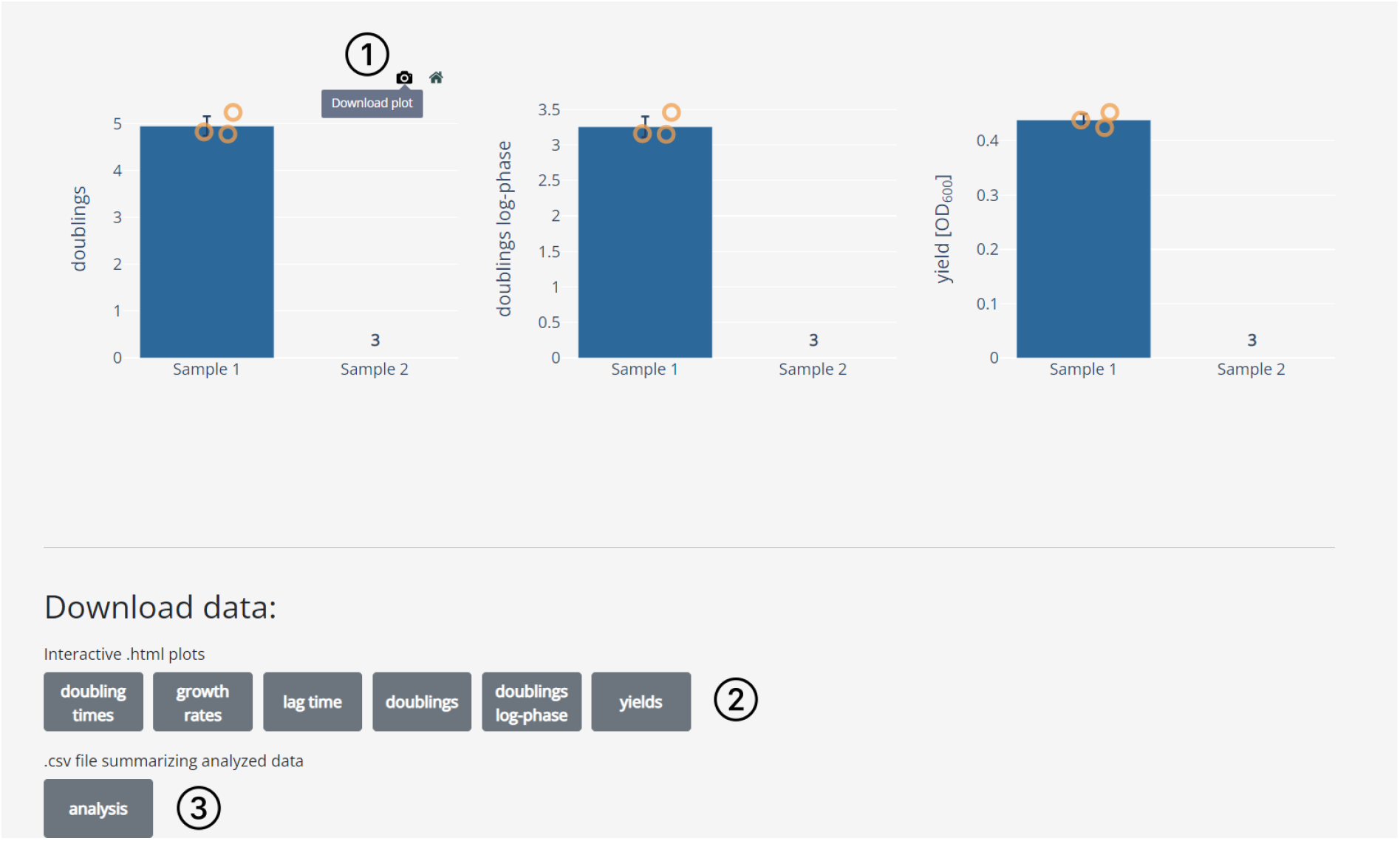
Dashing Growth Curves data download options. ① Hovering on the top right corner of every plot shows an option to download the respective graph as a vector graphic. ② The same graphs can also be downloaded as interactive HTML plots. ③ The extracted growth parameters for all samples can be downloaded as a comma separated values (.csv) file.

Lastly, Dashing Growth Curves handles hundreds to thousands of growth curves each with hundreds of time-resolved measurements (we tested datasets with up to half a million data points, i.e. the number of samples multiplied by the number of timepoints recorded per sample).

## Conclusions

Here, we introduce Dashing Growth Curves, an open-source web application to support researchers in analyzing growth curves quickly and reliably. Dashing Growth Curves requires no programming skills and is operating system independent. Lastly, it gives users the flexibility to fit their data with different growth models or to use the traditional approach of manual selection of the logarithmic growth phase.

## Availability and requirements

Project name: Dashing Growth Curves

Project home page: https://github.com/mretier/growthdash

Operating system: operating system independent

Programming language: Python 3.11

License: GNU General Public License v3.0

Any restrictions to use by non-academics: None

## Author contributions

M.A.R. conceived the project and developed the software. M.A.R. and J.A.V. wrote the manuscript.

## Acknowledgements

We thank Timothy Bradley and Thomas Gassler for critical reading of the manuscript and Timothy Bradley, Thomas Gassler and Lars Büchel for providing feedback for the application. This work was supported by ETH Zurich and a grant from the Swiss National Science Foundation (310030B-201265).

